# IL-6 Evades KSHV-Mediated Hyperadenylation repression via CRM1-Dependent Nuclear Export

**DOI:** 10.1101/2025.06.05.657986

**Authors:** Samantha Schultz, Jacob Miles, Will Dwyer, Daniel MacVeigh-Fierro, Mandy Muller

## Abstract

RNA turnover is critical for regulating cellular homeostasis, with nuclear export representing a key step in mRNA fate. During infection by Kaposi’s Sarcoma-associated Herpesvirus (KSHV), widespread mRNA decay is mediated by the viral endonuclease SOX, which depletes cytoplasmic transcripts and induces secondary nuclear RNA processing defects. One such defect includes transcript hyperadenylation, which promotes nuclear retention and decay. However, a subset of mRNAs escapes both SOX degradation and nuclear retention, raising questions about their export mechanisms. Here, we investigate how KSHV infection impacts mRNA poly(A) tail length and nuclear export dynamics using poly(A)-sequencing in KSHV-positive cells. Our data confirm a global increase in poly(A) tail length during KSHV infection, yet we identified a group of hyperadenylated transcripts that remain localized in the cytoplasm, suggesting active evasion of nuclear retention. Notably, we focused on interleukin-6 (IL-6), an mRNA known to escape SOX-mediated decay. Using G/I tailing and sPAT assays, we show that IL-6 is hyperadenylated yet, exported. We demonstrate that its export is dependent upon the CRM1 nuclear export pathway, rather than through the canonical NXF1-NXT1 pathway. Inhibition of CRM1 impairs IL-6 nuclear export and reduces steady-state mRNA levels, implicating CRM1 export in the stabilization of this transcript. Our findings reveal a previously unrecognized mechanism by which select host mRNAs like IL-6, bypass KSHV-imposed nuclear export block, thereby preserving their cytoplasmic function during infection. This study highlights viral manipulation of RNA processing and export pathways as a critical determinant of transcript fate and identifies CRM1 as a key mediator of selective transcript preservation during KSHV infection.

**SIGNIFICANCE STATEMENT:** Kaposi’s sarcoma-associated herpesvirus (KSHV) globally disrupts host gene expression by inducing host shutoff which triggers a global repression of the host transcriptome via widespread RNA decay, RNA, nascent transcript hyperadenylation and nuclear export blocks, yet a subset of transcripts escape this repression. This study reveals that despite acquiring long poly(A) tails, select host mRNAs such as IL-6 evade nuclear retention by using the alternative CRM1-dependent export pathway and remain stable in the cytoplasm. These findings challenge the prevailing model that hyperadenylation alone dictates nuclear decay and uncover a selective mechanism by which crucial host transcripts bypass KSHV-mediated gene repression. Understanding this selective escape provides new insights into host-virus interactions and highlights CRM1 as a potential therapeutic target in KSHV-associated diseases.

## INTRODUCTION

RNA turnover is a dynamic process that balances RNA synthesis and degradation, playing pivotal roles in regulating gene expression. Control over the stability of the transcriptome is critical to maintain cell homeostasis but also to respond to environmental stimuli, to adjust the cell response to stressors and to adapt expression in response to infections^1-3^. Yet, the factors that control RNA stability remain largely unknown. Many steps in mRNA processing occur in the nucleus and serve as landmarks for quality control pathways, ensuring that only viable transcripts are exported out of the nucleus^4^. mRNA export then unfolds in three main steps: recruitment of export factors, docking at the nuclear pore complex (NPC) and release in the cytoplasm. In recent years, it has emerged that mRNA export can be selectively regulated by specific export factors to control distinct biological processes^5^. The major export route for mRNAs relies on a heterodimer composed of NXF1 (nuclear export factor 1, also known as TAP) and NXT1 (NTF2-related export protein NXF1, also known as p15). The NXF1-NXT1 (or TAP-p15) heterodimer promotes docking at the NPC *via* direct binding to its mRNA cargo^6,7^. Additionally, RNA export *via* CRM1 (chromosome region maintenance 1, also known as exportin 1 or XPO1) is an additional export pathway, known to mediate nuclear export of select RNA, including miRNA, and mRNA subgroups^6,8-10^.

Given the critical importance of nuclear export, it is perhaps not surprising that viruses have evolved to extensively hijack nuclear export. The selective modulation of which transcripts reach the cytoplasm can grant the virus more access to translational resources and help dampen expression of antiviral genes^11-13^. This is exemplified by the fact that numerous viruses from Influenza A to Coronaviruses, Adenoviruses and Herpesviruses encode viral proteins that directly interfere with the nuclear export machinery^12,14-20^. An additional way virus can modulate nuclear export is to target steps of mRNA processing prior to export. One of the best example of this is during KSHV infection when it was shown that nuclear transcripts undergo hyperadenylation leading to their rapid nuclear decay^21,22^. This hyperadenylation occurs in the context of KSHV mediated *host shutoff*, a phenomenon initiated by the nuclease SOX, which promotes the cytoplasmic decay of host mRNAs^23-28^. This widespread cytoplasmic decay leads to long distance changes in the host cell, including release of RNA binding proteins from cytoplasmic targets, transcriptional shutdown and nuclear hyperadenylation^29-35^. Taken together, these combined viral-mediated attacks on the host transcriptome are particularly effective as it is estimated that up to 70% of the host transcriptome is downregulated during KSHV-mediated host shutoff^25,36,37^. This allows KSHV to thrive in its host cell, freeing translational resources that can now be re-directed towards viral transcripts and partially deactivating host defense systems.

However, one important gap remains in our understanding of KSHV takeover of nuclear export pathways during host shutoff: while most of the transcriptome is affected by SOX decay and the subsequent hyperadenylation phenotype, select transcripts escape this fate and reach the cytoplasm where they remain stable despite SOX nuclease activity^38-41^. We and others have shown that these spared transcripts play critical roles during infection and recent findings have uncovered important aspects of the mechanisms rendering these mRNAs refractory to SOX cleavage^40,42-44^. We thus set out to investigate whether these escaping mRNAs were also spared from viral-mediated hyperadenylation, and how these transcripts are exported out of the nucleus, despite the KSHV global nuclear export block.

Using high-throughput poly(A) sequencing, we show that hyperadenylation is prevalent during KSHV lytic infection, but a fraction of hyperadenylated mRNAs bypass the subsequent KSHV-induced nuclear export block and still reach the cytoplasm. We demonstrate that IL-6, the best characterized SOX cytoplasmic escapee acquires a longer tail during lytic infection which does not affect its stability. Furthermore, we show that the IL-6 transcript uses CRM1-dependent export to reach the cytoplasm rather than the canonical nuclear export *via* NXF1-NXT1, which is known to be blocked during KSHV infection. We therefore propose a model where select mRNAs, while not exempt from viral-mediated hyperadenylation, selectively escape KSHV-mediated nuclear block, and reach the cytoplasm where they also resist SOX-mediated RNA decay.

## RESULTS

KSHV infection broadly affects poly(A) tail length of host transcripts. To further characterize the global impact of KSHV on poly(A) length during infection we performed high-throughput RNA-sequencing on poly(A)-enriched RNA (Poly(A)-seq) purified from the KSHV positive cell line iSLK.WT either left latent or reactivated. To quantify the impact of poly(A) length on nuclear retention, cells were further fractionated to isolate RNA either from cell nuclei or cytoplasms. Using the transcription start site and transcription termination site as reference points, the distribution of reads was analyzed and showed that, as expected in poly(A)-seq, reads were enriched near 3’ end transcription termination sites and stop codons. We mapped the length of the poly(A) tail of 6969 unique genes across experimental conditions, of which 89% were protein coding genes (**Supplementary table 1**). The median length of poly(A) tails in our dataset was shifted up between our latent and lytic samples (length of 75 in latent cells and 83 in lytic cells) and as expected we observed an accumulation of longer poly(A) tails during KSHV lytic cycle as expected (**Figure 1A and Supplementary Figure 1**). However, RNAs harvested from the cytoplasmic fraction had much fewer long tails as shown by their length distribution (**Figure 1B**). We next focused on transcripts that had a poly(A) tail length that underwent notable modifications during KSHV lytic cycle. We found that 160 transcripts in the nucleus had significantly different tail lengths upon KSHV reactivation of which 97 had acquired longer tails. The dynamism of tail length appeared even more robust in the cytoplasm during KSHV lytic cycle, with 405 unique transcripts having significant changes to their tail length including 231 having much shorter tails, in line with figure 1B (**Figure 1C**). Intriguingly, and despite the current model of viral-induced hyperadenylation being a strict indicator of nuclear retention and nuclear decay, our results also indicate that 174 cytoplasmic transcripts appear to have longer poly(A) tails upon KSHV reactivation. This suggested to us that these newly hyper-adenylated transcripts successfully evaded nuclear retention and decay despite acquiring a longer poly(A) tail. We thus conducted a GO-term analysis on this pool of cytoplasmic transcripts with different poly(A) tail size during KSHV lytic infection (**Figure 1D**). Since poly(A) tail size in the cytoplasm is a predictor of stability, we reasoned that enrichment of certain GO-term based within transcripts with shorter poly(A) could indicate that these functions are downregulated and reversely, function enriched in hyper-adenylated transcripts could reveal potential upregulation of these pathways. In the pool of transcripts with longer poly(A) tails, we identified an enrichment in positive regulation of DNA-templated transcription, which could greatly benefit viral replication. Reversely, in the transcripts that present with shorter poly(A) tails, we found an enrichment of mitochondrial electron transport which could indicate that important regulators of cell homeostasis are targeted by KSHV.

**Figure 1:**
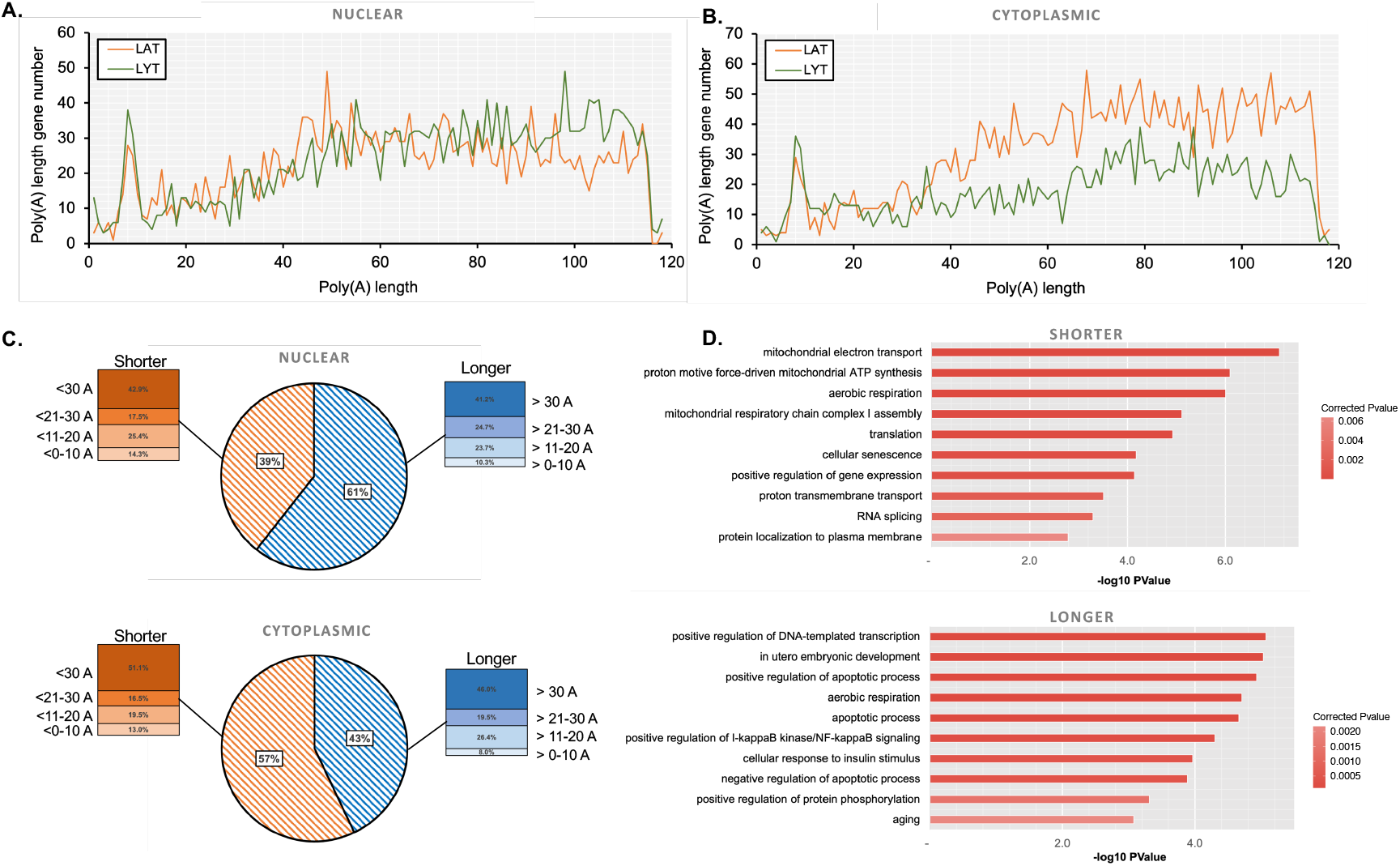
Poly(A)-seq reveals dynamic regulation of poly(A) tail size during KSHV lytic infection. The KSHV positive cell line iSLK.WT was either left latent or reactivated. Cells were then fractionated, and RNA harvested from the nuclear or cytoplasmic compartment. Poly(A)-seq library preparation and data analysis were performed as described in the method section. Nuclear (**A**) or Cytoplasmic (**B**) extracted mRNA poly(A) tail length were measured. The graphs represent the distribution of poly(A) length across mapped reads. **C**. Proportion of shorter (orange) or longer (blue) size tail within the nuclear (top) or cytoplasmic (bottom) pool of transcript that have differential tail lengths. The zoom in section shows the difference within these categories with tail shortening or lengthening between 0 and 30 As. D. Go-term analysis using AmiGO to determine the top 10 most enrichment “processes” with the transcripts with shorter (top) or longer (bottom) polyA tails in the cytoplasm.

**Figure 2:**
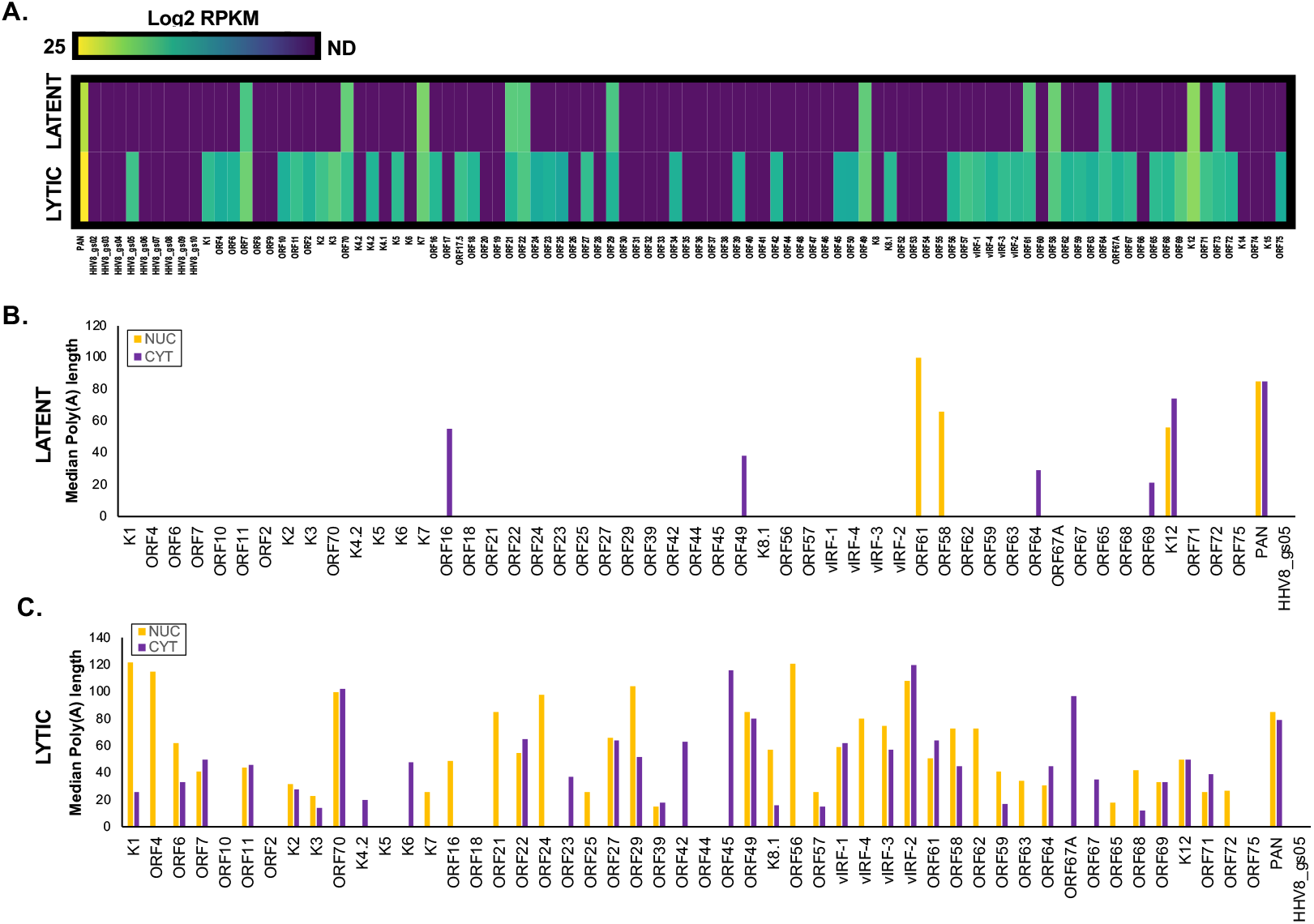
Poly(A) tail size of KSHV transcripts does not affect cytoplasmic expression. A. The KSHV positive cell line iSLK.WT was either left latent or reactivated. Cells were then fractionated, and RNA harvested from the nuclear or cytoplasmic compartment. Expression was determined based on RPKM reads and plotted as log2 RPKM. ND refers to <u>N</u>on-<u>D</u>etected transcripts. **B-C:** Poly(A)-seq library preparation and data analysis were performed as described in the method section. Nuclear or Cytoplasmic extracted mRNA poly(A) tail length were measured. The graphs represent the median poly(A) length across mapped reads in latent (B) or lytic cells (C).

Of note, given that samples were harvested in the KSHV-positive cell line iSLK.WT, we also detected highly expressed viral genes across our latent and lytic samples, allowing us to map the poly(A) tail length of 53 KSHV genes (**Supplementary table 2 and Figure 2**). As opposed to host genes, poly(A) tails on viral transcripts -when detected both in the nucleus and the cytoplasm-were consistent in size, suggesting a less dynamic process. For the few viral transcripts detected both in latent and lytic samples, we also observed that their poly(A) tails were not elongated, possibly suggesting that these viral transcripts are not susceptible to the global hyperadenylation event in the nucleus or are escaping the nuclear export block quickly enough to evade it. Overall, our poly(A)-seq data confirms that there is a global hyperadenylation of host transcripts during KSHV lytic infection but also reveal that certain transcripts may still reach the cytoplasm carrying longer poly(A) tails, suggesting a possible escape mechanism from nuclear retention and decay.

### IL-6, a known SOX escapee, is hyperadenylated and exported to the cytoplasm

We have previously shown that select host transcripts are refractory to SOX cleavage in the cytoplasm^38-40^. Given that these resistant mRNAs are readily expressed in the cytoplasm and remain stable in the face of SOX-induced decay, we hypothesized that these host transcripts must also be resistant to viral-mediated hyperadenylation and subsequent nuclear decay or at least evade the nuclear retention phenotype caused by this hyperadenylation. We therefore next wanted to assess whether these SOX-resistant transcripts where also hyperadenylation resistant. To do so, we focused on the host transcript IL-6 which is the best characterized SOX escapee and plays a critical role for KSHV-associated pathogenesis. We first fractionated iSLK.WT cells to verify the localization of IL-6 mRNA prior to host shutoff and subsequent hyperadenylation. Using two non-coding predominantly nuclear RNA as controls (Xist and Malat1), we observed that in latent cells, IL-6 mRNA is readily detectable in cytoplasmic fractions as expected (Figure 3A). Surprisingly, in lytically reactivated cells, we observed that the ability of IL-6 mRNA to be exported to the cytoplasm was preserved. Therefore, we next wanted to assess whether IL-6 mRNA was spared from host shutoff-associated hyperadenylation. To do so, we performed G/I tailing on RNA extracted from nuclear and cytoplasmic fractions in cells either left latent or reactivated. As a control, we used GAPDH mRNA as this mRNA was previously shown to be unaffected by hyperadenylation in the context of KSHV lytic cycle^16^. As shown in Figure 3B both IL-6 and GAPDH mRNA are present in nuclear and cytoplasmic fractions. However, contrary to the constant poly(A) tail size of GAPDH across conditions, we found that IL-6 tail was shifted up upon KSHV lytic reactivation. To confirm this result, we next measured IL-6 poly(A) tail profile using sPAT (splint-mediated PolyA Test^45^) to better capture the hyperadenylation pattern on IL-6 tail. We used two amplification approaches: one where we used an IL-6 primer specific annealing temperature which predominantly amplified one size of poly(A) tail, likely representative of the most abundant average size; and one where we used a universal annealing temperature, less optimized for our IL-6 primer but allowing the capture of poly(A) “smears”. Here again, we confirmed that IL-6 mRNA appears to be hyperadenylated during KSHV lytic cycle (**Figure 3C**). Taken together, these results suggest that contrary to our initial assumption, IL-6 does not evade host shutoff-associated hyperadenylation yet appears to readily evade nuclear retention and continue to be exported out of the nucleus.

**Figure 3:**
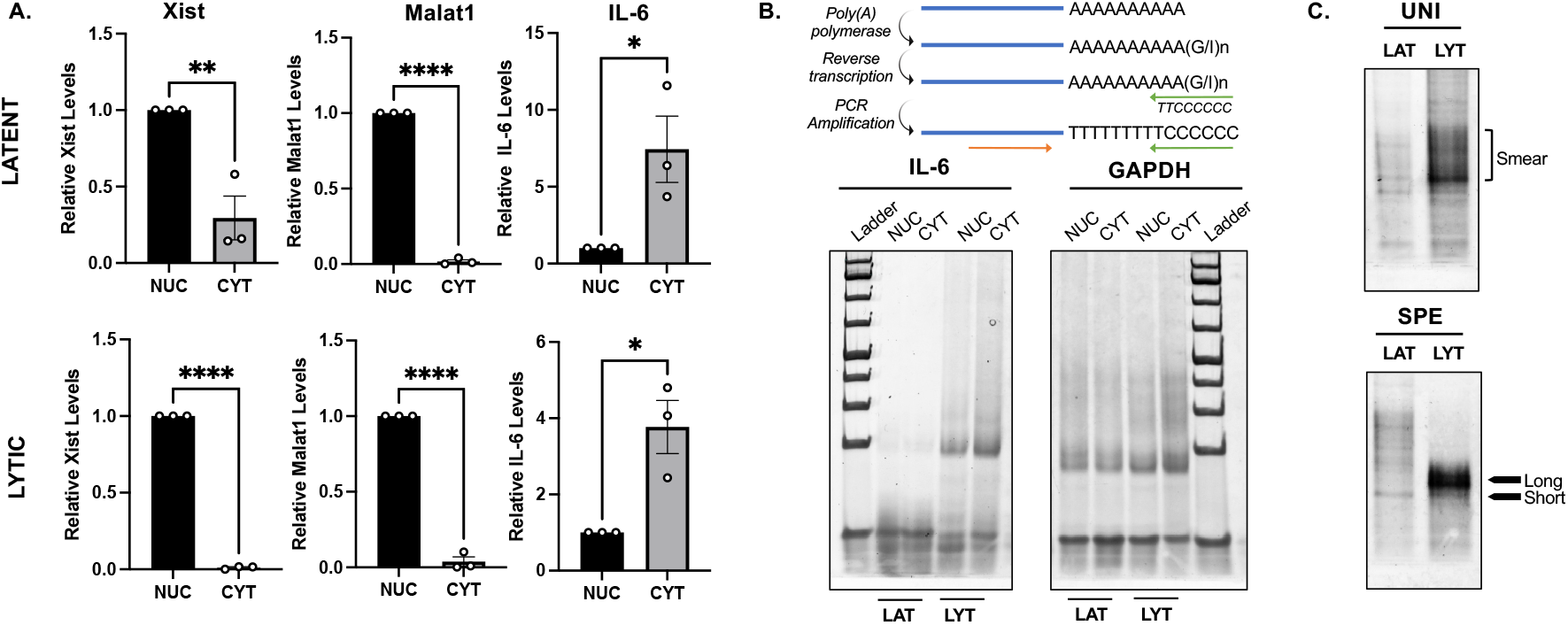
IL-6 is hyperadenylated yet still reaches the host cell cytoplasm. **A**. iSLK.WT cells were either left latent (Top) or reactivated (bottom) then fractionated, and RNA extracted from nuclear (NUC) or cytoplasmic (CYT) fractions. RNA was subjected to RT-qPCR with the indicated primers. **B**. RNA extracted as described in A was subjected to G/I tailing followed by reverse transcription and PCR amplification using either IL-6 (left) or GAPDH (right) specific primers according to the Affymetrix kit. PCR products were then run on a 2% agarose gel and stained with SYBR gold. **C**. Total RNA was extracted from iSLK.WT cells either left latent (LAT) or reactivated (LYT) and used for sPAT using either a universal annealing temperature of 60°C for the final amplification (UNI – Top) or the IL-6 primer specific annealing temperature (SPE – Bottom). PCR products were then run on a Criterion 5% TBE-Agarose Gel and stained with SYBR Gold.

### IL-6 transcript uses CRM1 to shuttle out of the nucleus

Given that during KSHV-mediated host shutoff, hyperadenylated transcripts are believed to be retained in the nucleus, we next wanted to assess how IL-6 is exported. Since it has been previously shown that the NFX-TAP export pathway is impaired upon host shutoff^35^, we thus reasoned that IL-6 could use an alternative nuclear export route to reach the cytoplasm. We thus treated cells with a CRM1 inhibitor Eltanexor (KPT8602) or DMSO as vehicle control and fractionated our iSLK.WT cells either left latent or reactivated, allowing us to measure the ability of IL-6 to reach the cytoplasm when CRM1 is inhibited. As shown in **figure 4A**, IL-6 mRNA levels are significantly lower in the cytoplasm upon Eltanexor treatment. Intriguingly, this is true both in latent and lytic cells, suggesting that IL-6 commonly uses this export pathway, not solely in response to its hyperadenylation. We next wondered whether IL-6 nuclear retention upon CRM1 inhibition would affect IL-6 stability. We thus extracted total RNA from cells treated with Eltanexor (or DMSO as vehicle control) and measured IL-6 steady state levels by RT-qPCR. As predicted, IL-6 steady state levels decrease upon CRM1 inhibition (**Figure 4B**), suggesting that its ability to escape nuclear retention allows it to evade hyperadenylation-associated nuclear decay.

**Figure 4:**
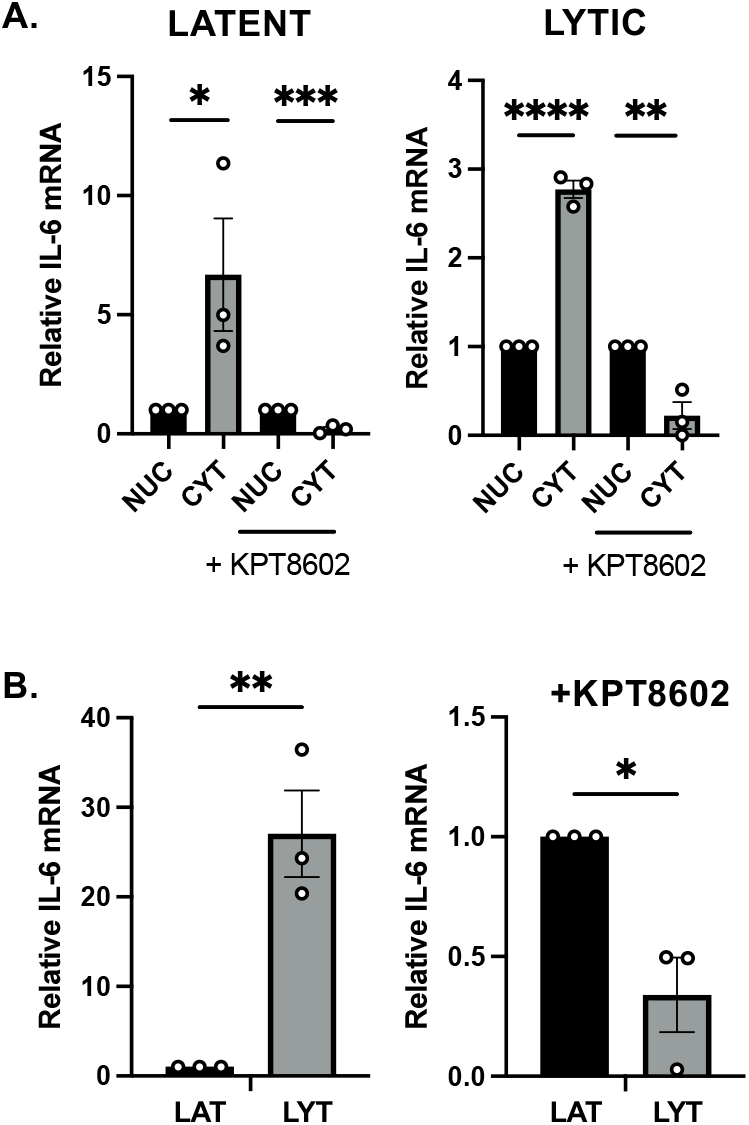
IL-6 mRNA relies on CRM1 for nuclear export and stability. iSLK.WT were treated with a CRM1 inhibitor KPT8602 (or DMSO as control) and either left latent (left) or reactivated (right). **A**. Cells were then fractionated, and RNA harvested from nuclear (NUC) or cytoplasmic (CYT) fractions and used for RT-qPCR. **B**. RNA was extracted from whole cells and used to measure IL-6 steady state levels by RT-qPCR.

## DISCUSSION

The ability to selectively regulate host gene expression is critical for successful viral infection and viruses have evolved diverse and refined strategies to manipulate RNA fate. This is particularly evident during Kaposi’s sarcoma-associated herpesvirus (KSHV) infection, where the viral endonuclease SOX induces an extensive host shutoff program. This leads to widespread cytoplasmic RNA decay, transcriptional repression, hyperadenylation, and nuclear retention of host transcripts, which together impose strict control over which host genes are expressed and enhancing viral gene expression. However, we and others have previously shown that a subset of host transcripts can evade SOX-mediated cytoplasmic degradation. This prompted us to ask whether these same transcripts also escape other layers of repression, particularly the aberrant nuclear hyperadenylation associated with KSHV lytic infection. Given the well-established role of 3′ end processing in determining mRNA stability and export, we initially hypothesized that SOX-resistant transcripts would also evade hyperadenylation. To test this, we globally profiled poly(A) tail lengths in cells undergoing KSHV lytic reactivation. Consistent with prior observations, we found a global increase in poly(A) tail length following reactivation. However, our data revealed a more nuanced picture: while nuclear transcripts tended to show elongated poly(A) tails, many cytoplasmic transcripts also exhibited longer poly(A) tails. This suggests that hyperadenylation alone cannot fully explain nuclear retention during KSHV infection. The poly(A) tail length of cytoplasmic transcripts plays a crucial role in determining both their stability and translational rates; therefore, it is important to better characterize how acquiring longer poly(A) tails may shape cytoplasmic regulation. Nonetheless, these transcripts, once exported, still face SOX-mediated degradation, indicating that surviving host transcripts must overcome both nuclear and cytoplasmic barriers.

Interestingly, viral transcripts were largely unaffected by changes in poly(A) tail length. Of the few viral RNAs detectable in both latent and lytic states, none showed significant poly(A) length alterations upon reactivation. Similarly, viral RNAs found in both nuclear and cytoplasmic compartments maintained consistent tail lengths. This may be explained by the activity of viral proteins such as ORF10, which facilitate efficient export of viral transcripts despite a general shutdown of host nuclear export pathways. Therefore, it is likely that viral transcripts transit rapidly through the nucleus, thereby avoiding the repressive effects of hyperadenylation.

Given the complex repression landscape faced by host mRNAs during KSHV lytic infection, we focused our attention on IL-6, a host transcript known to evade SOX-mediated mRNA decay. We initially hypothesized that IL-6 might escape nuclear repression by avoiding hyperadenylation. Surprisingly, however, IL-6 mRNA exhibited an elongated poly(A) tails during lytic infection. To understand how IL-6 bypasses nuclear retention induced by hyperadenylation, we identified CRM1 (also known as exportin 1 or XPO1)-dependent export as essential for its nuclear export. Intriguingly, IL-6 relies on CRM1-mediated export in both latent and lytic cells, suggesting this is a constitutive pathway rather than an adaptation to its host shutoff-associated hyperadenylation. CRM1-dependent nuclear export typically requires adaptor proteins to bridge RNA cargo and the export machinery. We have previously characterized the ribonucleoprotein complexes associated with SOX-resistant transcripts such as IL-6. Future studies should investigate whether any of these RNA-binding proteins function as CRM1 adaptors for IL-6 or other escapee transcripts. Furthermore, recent studies have identified CRM1 as a critical regulator in KSHV-transformed cells and in the host immune response to KSHV replication^46,47^. Our findings add to this body of evidence by highlighting CRM1’s role in preserving the expression of select host transcripts during viral infection, further supporting its potential as a therapeutic target in KSHV-associated disease.

In summary, our results reveal a dynamic and complex regulation of poly(A) tail length during KSHV lytic infection. Given the central role of poly(A) tails in RNA fate determination, it is not surprising that KSHV extensively manipulates this process. Nonetheless, select host mRNAs such as IL-6 appear capable of resisting both nuclear retention and cytoplasmic degradation despite being susceptible to hyperadenylation. IL-6 has long been implicated in KSHV biology, and its persistent expression during host shutoff reinforces its importance as a key factor in viral pathogenesis.

## Materials and Methods

### Cells and transfections

The KSHV-infected renal carcinoma human cell line iSLK.BAC16 (iSLK.WT) bearing a doxycycline-inducible RTA were grown in Dulbecco’s modified Eagle’s medium (DMEM; Invitrogen) supplemented with 10% fetal bovine serum (FBS). KSHV Lytic reactivation was induced by the addition of 1 μg/ml doxycycline (BD Biosciences) and 1 mM sodium butyrate for 48hr. For DNA transfections, cells were plated and transfected after 24h when 70% confluent using PolyJet (SignaGen). CRM1 inhibitor KPT-8602 (Eltanexor) was diluted in DMSO and used at a concentration of 0.5 uM for 1h prior to lytic reactivation as previously^47^.

### RT-PCR

Total RNA was harvested using TRIzol according to the manufacturer’s protocol. cDNAs were synthesized from 1 μg of total RNA using AMV reverse transcriptase (Promega) and used directly for quantitative PCR (qPCR) analysis with the SYBR green qPCR kit (Bio-Rad). Signals obtained by qPCR were normalized to 18S unless otherwise noted. Primers used in this study are as follows: IL-6 [Fw: ACTCACCTCTTCAGAACGAATTG; Rv:CCATCTTTGGAAGGTTCAGGTTG], Malat1 [Fw: GAATTGCGTCATTTAAAGCCTAGTT; Rv:GTTTCATCCTACCACTCCCAATTAAT], Xist [Fw: GTAGGTGTGCTGATAACCAAGGC;Rv: GGGAAAGGAAGATTGAGGGTGG].

### G/I tailing

Cytoplasmic and nuclear fractions of total RNA were prepared 48h after reactivation in iSLK.WT cells as described before^48^. Briefly, cell pellets were resuspended in 200ul of lysis buffer (50 mM Tris, pH 8.0, 140 mM NaCl, 1.5 mM MgCl2, 0.2% Nonidet P40, 1 mM DTT, and 200 units of RNAsin), incubated 5 min on ice, and spun 2 min at 3000g. Cytoplasmic RNA was extracted from the supernatant while nuclear RNA was collected from the pellet using TRIzol according to the manufacturer’s protocol. G/I tailing was performed using the USB Poly(A) Tail-Length Assay Kit (affymetrix) using the kit’s universal reverse primer and the gene specific primers as follows: IL-6 [CCAGATCATTTCTTGGAAAGTGTA] and GAPDH [TGAATCTCCCCTCCTCACAGTT]. Samples are then run on a 5% TBE acrylamide gel and stained with SYBR gold.

### sPAT

Total RNA was extracted by TRIzol according to the manufacturer’s protocol. The RNA anchor [CAGCUGUAGCUAUGCGCACCGAGUCAGAUCAG] was ligated using splint ligation and after DNAse treatment, the samples were phenol-chloroform extracted and reverse transcribed using the following primer: Fw: CTGATCTGACTCGGTGCGCA Rv: TGCGCATAGCTACAGCTGTTTT. Reverse transcription was run either using a universal annealing temperature of 60C (referred to as UNI) or at the specific annealing temperature recommended for the primers used (referred to as SPE). Samples were then run on a 5% TBE acrylamide gel and stained with SYBR gold.

### PolyA sequencing

To prepare the samples for sequencing, total RNA was fractionated using the Norgen Cytoplasmic & Nuclear RNA Purification Kit. After extraction and precipitation, samples were sent to CD genomics for further processing. Total RNA was treated with RQ1 DNase (promega) to remove DNA. The quality and quantity of the purified RNA were determined by measuring the absorbance at 260 nm/280 nm (A260/A280) using smartspec plus (BioRad). RNA integrity was further verified by 1.5% agarose gel electrophoresis. For each sample, 5μg of total RNA was used for poly(A)-seq library preparation. Total RNA was digested by RNase T1 (Thermo, EN0541), polyadenylated mRNAs were purified and concentrated with oligo(dT)-conjugated magnetic beads (Vazyme, N401-01) before directional poly(A)-seq library preparation. After 3’ adaptor (Gnomegen, K02420-L) ligation, purified mRNAs were reverse transcribed to cDNA with RT primer that was complemented with the 3’ adaptor. Then DNA was synthesized with Terminal-Tagging oligo cDNA and SMARTer® Stranded RNA-Seq Kit (TAKARA, 634837). For high-throughput sequencing, the libraries were prepared following the manufacturer’s instructions and applied to Novaseq Xplus for 150nt paired-end sequencing. During library preparation, poly(A) tail spike-in sequences of lengths 120nt and 40nt were added as controls. Raw reads were filtered to get high-quality data and processed as follows: poly(A) sequences were first searched in the reads from the right side, with a length of 6A and a mismatch rate of 0.2. followed by search for poly(A) sequences from the left side, with a length of 9A and a mismatch rate of 0.1. Reads containing 4 consecutive non-A bases were truncated keeping the longest portion of the poly(A) sequence. If the remaining part of the reads after truncating the poly(A) sequence has a length of at least 20, then such reads are classified as poly(A) clean reads, and the corresponding poly(A) length is equal to the length of poly(A) in the reads. The retained poly(A) sequence lengths are greater than or equal to 11 nt, and the alignment was performed using Bowtie2 against the human genome (GRCh38.p13) as well as the KSHV genome (RefSeq: GCF_000838265.1). Expression was normalized for mRNA length and sequencing depth (RPKM - Reads per kilobase of a gene per million reads), ensuring the expression abundance between different sequencing samples can be compared. The raw sequence data generated in this paper were deposited on the NCBI Gene Expression Omnibus (GEO) server under the identifier GSE298531.

### Statistical analysis

All results are expressed as means ± standard errors of the means (SEMs) of experiments independently repeated at least three times. Unpaired Student’s t test was used to evaluate the statistical difference between samples where indicated. For non-gaussian distribution, the Wilcoxon text was used. Significance was evaluated with P values as follows: * p<0.05; ** p<0.01; *** p<0.001.

## SUPPORTING INFORMATION

**Supplementary Figure 1:** median poly(A) distribution across the host transcriptome represented as a violin plot with individual values.

**Supplementary Table 1:** Poly(A) tail length of each unique genes averaged over replicate data in each condition (latent vs. lytic; Nuclear vs. cytoplasmic fractions).

**Supplementary Table 2:** Poly(A) tail length of each unique viral genes averaged over replicate data in each condition (latent vs. lytic; Nuclear vs. cytoplasmic fractions).

## Acknowledgments

We would like to thank all the members of the Muller lab for their helpful insights. The genomics services and analyses were performed by CD genomics.

## Data Availability

The RNA-seq data was deposited in NCBI Gene Expression Omnibus (GEO) database under accession numbers GSE298531.

## Funding Statement

M.M. was supported by NIH grant R35GM138043.

## Notes

### Competing Interest Statement

The authors have declared no competing interest.

