## Supplementary figures and images for "IL-6 Evades KSHV-Mediated Hyperadenylation repression via CRM1-Dependent Nuclear Export"

### supplementary figure 1

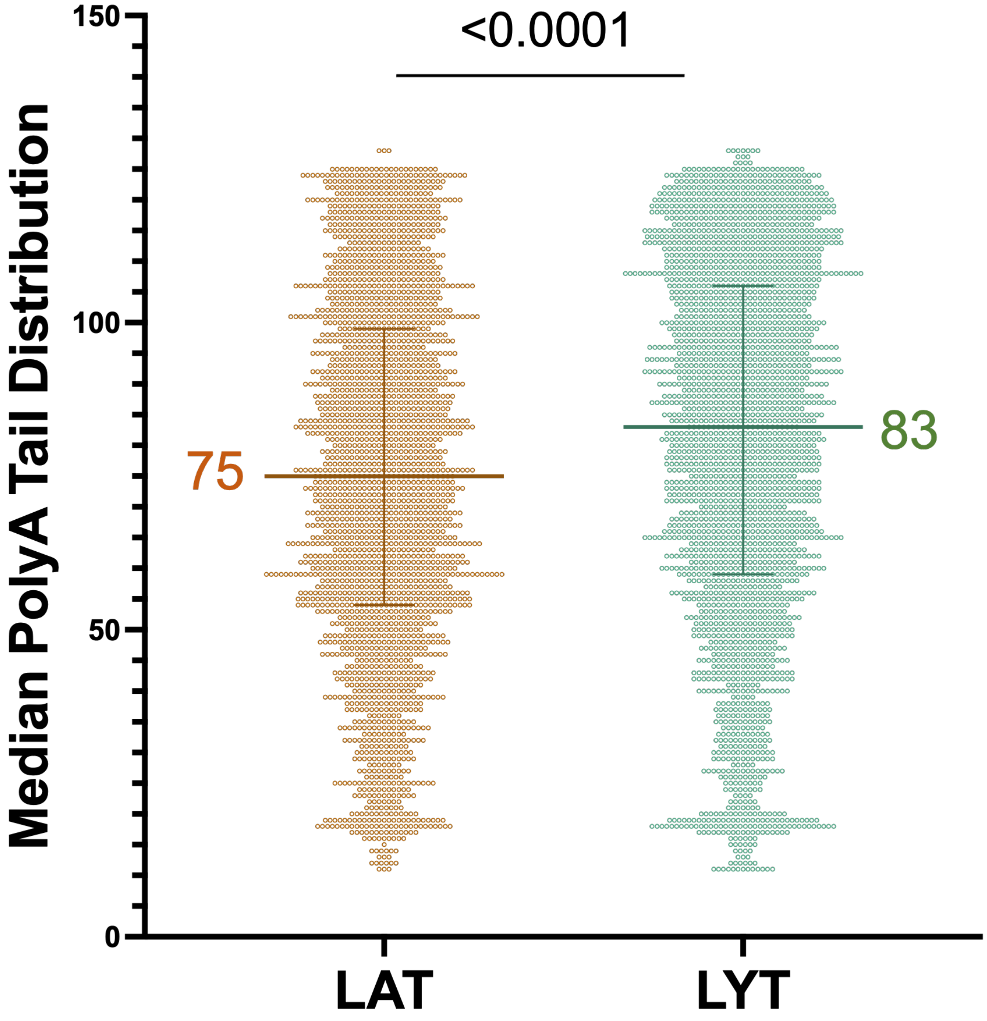
